# The scaling behavior of hippocampal activity in sleep/rest predicts spatial memory performance

**DOI:** 10.64898/2026.03.11.711043

**Authors:** Predrag Živadinović, Fabrizio Lombardi, David Dupret, Charlotte Boccara, Sofia Taveira, Juan Ramirez-Villegas, Gašper Tkačik, Jozsef Csicsvari

## Abstract

In the hippocampus, reactivation of spatial memories during sleep enhances consolidation and recall. However, network dynamics may independently influence memory retention. We applied the Phenomenological Renormalization Group to CA1 neuronal activity in rats during sleep/rest epochs surrounding a spatial learning task. The scaling exponent of activity variance (***α***), assessed either before or after learning, predicted subsequent recall performance independently of reactivation. The prediction model was transferable across subjects, suggesting that it is a robust biomarker of learning ability. Single-cell features, such as burst propensity and intrinsic timescales, correlated with ***α*** and memory, ***α*** predicted retention even when controlling for these factors. Our results identified a link between scale-free circuit dynamics and memory stabilization, suggesting that the tuning of the underlying dynamical regime near criticality is a key determinant of memory longevity. Targeted modulation of these collective states could offer new avenues for memory enhancement.

## Introduction

Decades of research have established sleep as a critical window for memory stabilization, supporting the long-standing hypothesis that systems consolidation primarily occurs during offline states(*1–6*). At the circuit level, the selective reactivation of neural ensembles during sleep has provided a key mechanistic framework for how these processes unfold (*7–11*). However, it remains unknown whether the underlying dynamical state of the hippocampal circuit exerts an independent, overarching influence on the efficiency of consolidation and, ultimately, memory longevity. Establishing such a link would demonstrate that stabilization is governed by global network dynamics, opening new avenues for memory enhancement.

The capacity of neural circuits to balance stability and flexibility of their activity states is central to their function. Dynamical systems theory suggests that brain functions should be optimized near the transition between sub- and supercritical neural avalanche regimes (*12–15*); these transitions often coincide with the emergence of neural oscillations (*16*). While demonstrating such tuning *in vivo* remains an experimental challenge, analytical signatures of criticality, such as scale-free patterns and power-law distributions have been identified (*17–20*). These can vary across behavioral states and correlate with sensory discrimination (*21, 22*).

The hippocampus is a key structure for navigation and spatial learning (*23*). While place cells there encode the animal’s location, the reactivation of their activity patterns during sharp-wave ripples (SWRs) (*24*) predicts the accuracy of subsequent spatial memory recall (*8*). This implies that the frequency and fidelity of reactivation are the primary determinants of learning efficiency (*10, 11, 25, 26*). What remains unknown, however, is whether collective network dynamics influence memory stabilization beyond, and independently of, specific ensemble reactivation.

Here, we tested the hypothesis that recall after sleep/rest is governed by hippocampal circuit dynamical regimes characterized by their scale-invariant properties. Using the Phenomenological Renormalization Group (PRG) (*18*), we identified scale-free features of network dynamics and found that the scaling of activity variance predicted memory retention across animals. Our results suggest that the underlying circuit dynamics associated with a near-critical, scale-free regime provide an essential substrate for memory longevity.

## Results

We analyzed CA1 pyramidal cell and interneuron activity from a cheeseboard learning task (Fig. 1A), using published data: (*8*), n=3 rats, 7 recording days; (*27*) n=2 rats, 5 days; and new recordings, n=6 rats, 10 days. Animals learned three reward locations daily, followed by a memory retention test (post-probe) after a period of sleep or rest (≳30 min, post-learning rest). Baseline controls included pre-learning probe (pre-probe) and sleep/rest sessions. In the post-probe session, animals spent significantly more time in the vicinity of reward locations compared to the pre-probe baseline, confirming successful memory retention across the intervening rest period (Fig. 1B).

**Figure 1.**
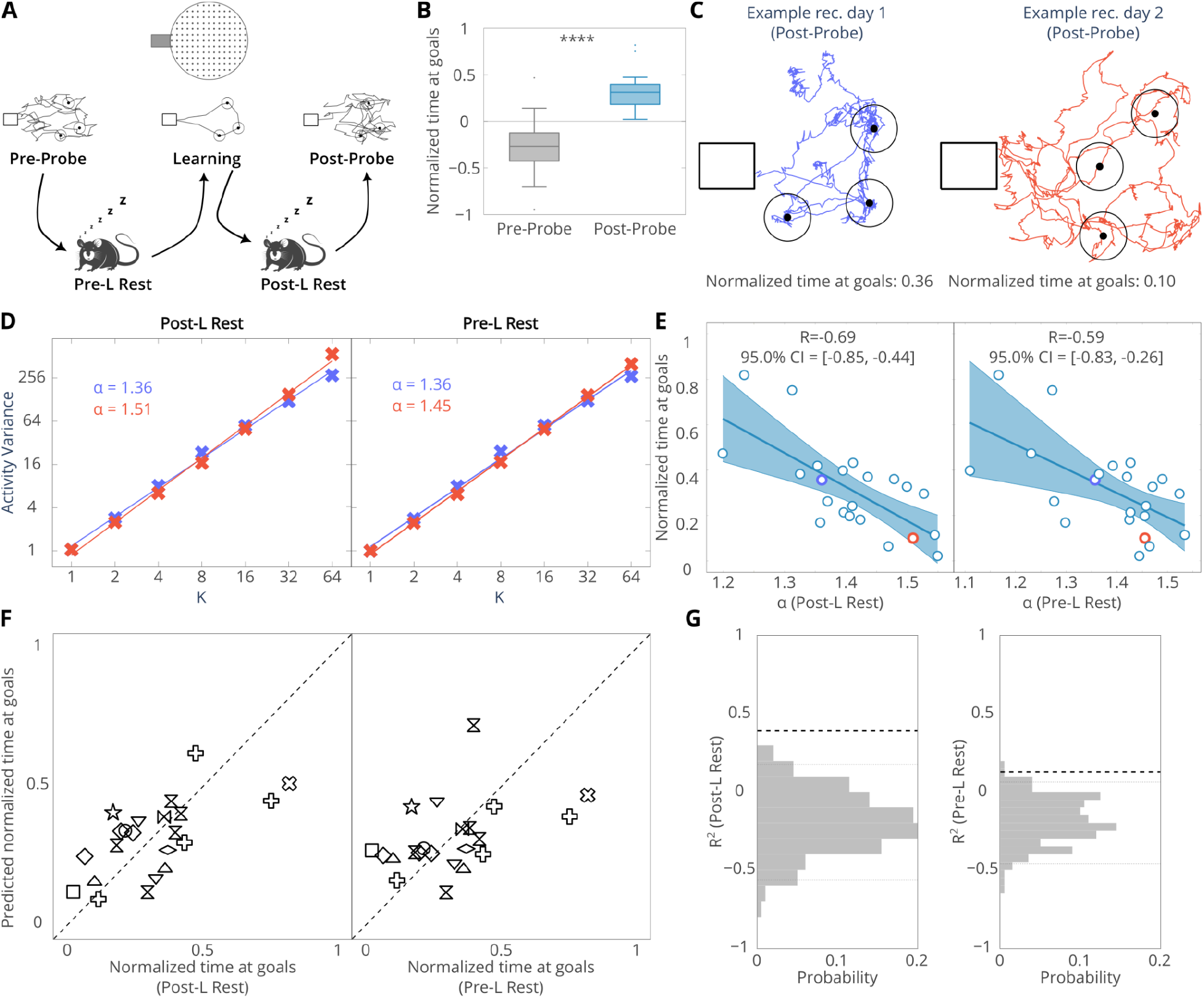
Scaling of activity variance predicts spatial memory performance. (**A**) Top: Sessions of the cheeseboard paradigm. Middle: Rats were trained to locate hidden food rewards at three goal locations (circles); sample trajectories with start box (rectangle) are depicted. Bottom: Learning was preceded and followed by sleep/rest (Pre-L Rest, Post-L Rest) and memory was compared on unrewarded Pre-Probe (control) and Post-Probe sessions. (**B**) Normalised time at goals is higher in the post-probe compared to the pre-probe (p < 0.0001, Wilcoxon signed-rank test). (**C**) Post-probe sessions from two representative recording days, showing good (blue) and weaker (red) recall. (**D**) Scaling of activity variance (***α***) during post-learning (left) and pre-learning rest (right), from the sessions in (C). Lower ***α*** implies better recall. (**E**) Normalized time at goals in post-probe correlates with ***α*** in post-learning (left) and pre-learning rest (right); color corresponds to sessions in (C, D). (**F**) Memory retention prediction from ***α*** across animals (different plot symbols; each animal’s retention was predicted with a model fitted using the other animals), ***α*** measured either in post-learning (left) or pre-learning rest (right). (**G**) Coefficient of determination for prediction models in (F); dashed black line: true model; gray histograms: shuffled models; dashed grey lines: 95% CI.

To characterize hippocampal dynamics across scales of activity, we applied the PRG (*18*) to the recorded multiple units in CA1 (Fig. 2). The PRG method employs an iterative coarse-graining procedure that simplifies network activity by successively merging units based on their correlation structure. In this procedure, two units with the highest pairwise correlation are combined first, followed by merging the highest-correlated pair among the remaining units; the procedure repeats until all units are merged. Each step effectively halves the total number of units, while the number of original neurons within each coarse-grained unit increases exponentially: K=2^k^ at step k (Fig. 2A). This iterative process reveals whether the activity and correlation structures remain self-similar across scales (Fig. 2B, 2C).

**Figure 2.**
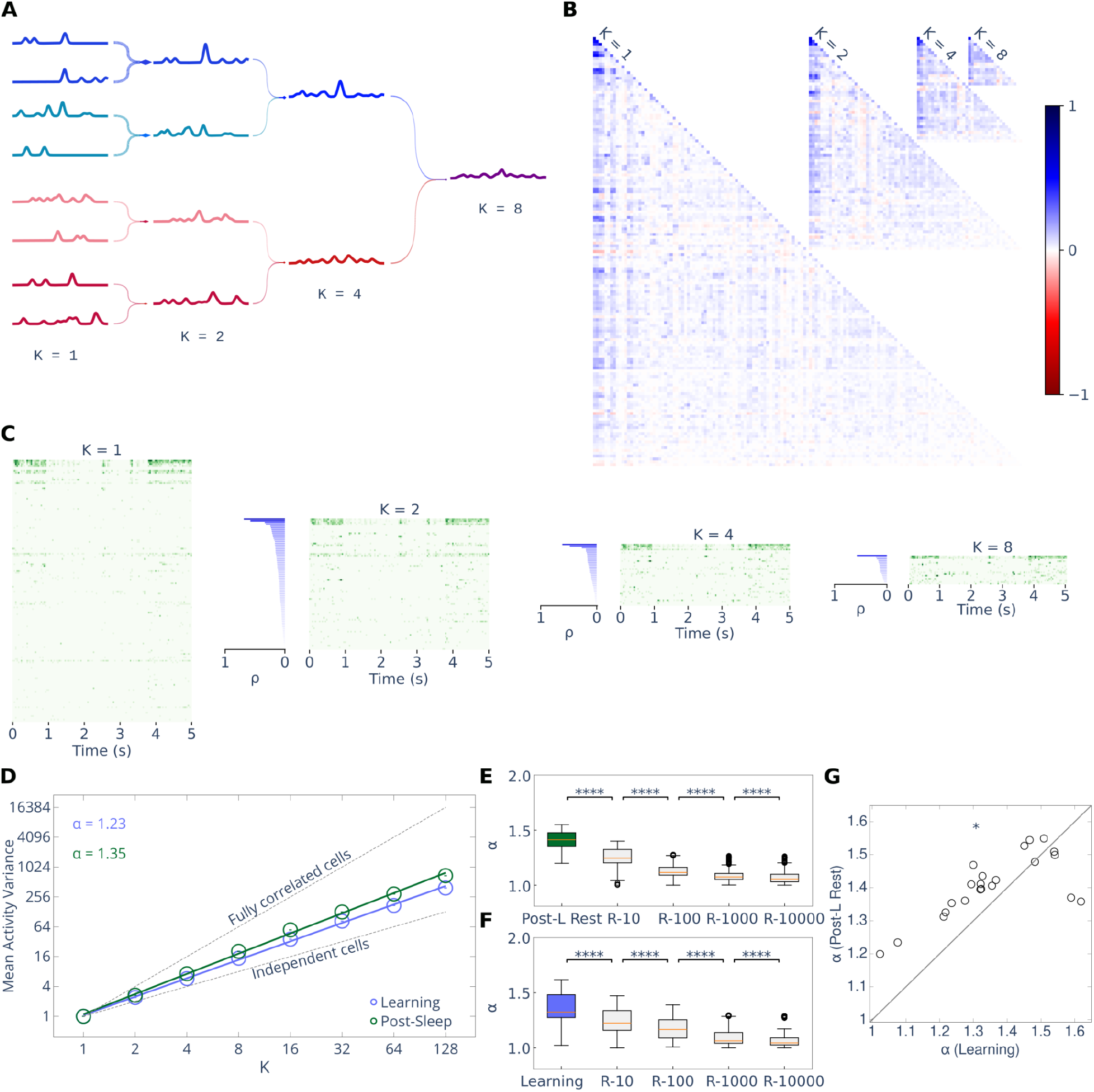
Phenomenological Renormalization Group for hippocampal activity. (**A**) Illustration of the PRG. Each line represents the normalised activity of a unit over time. Pairs of units with the strongest correlation are merged (line colors illustrate the correlation strength of merged pairs); the summed activity is normalised. K denotes the number of cells in each coarse-grained unit. (**B**) Pairwise correlation matrices for different K in one sample session. (**C**) Activity matrices (in green) for cells in (B);blue: corresponding pairwise correlation distribution. (**D**) Scaling of activity variance (***α***) during Learning (blue) and Post-L Rest (green). Same session as in (B,C). (**E, F**) Scaling exponent ***α*** in Post-L Rest (E) and Learning (F) vs. randomised controls (see Methods) using increasing time window (**** p < 0.0001 in two-sided t-test). (**G**) ***α*** is higher in post-learning rest than in learning (p < 0.05, Wilcoxon signed-rank test).

At each coarse-graining step, we measured network activity properties to determine if they scale with K (Fig. 2D, S1). Specifically, the scaling exponent ***α*** quantifies how the average variance of coarse-grained unit activity scales with the scale K on a log-log plot. The ***α*** characterises the degree of coordination between individual neurons; ***α*** = 1 indicates completely uncorrelated activity, whereas ***α*** = 2 represents a fully correlated neural population. In our recordings, ***α*** values indicated partially correlated dynamics. When we disrupted these correlations by jittering or randomising spikes across various timescales (see Methods), ***α*** progressively decreased (Fig. 2E-F, S2A-B), suggesting that neuronal correlations exert a long-range temporal influence extending to tens of seconds.

We found that ***α*** depended on the behavioral state, with significantly higher values in post-learning rest than during the learning task (Fig. 2G), a difference not observed in pre-learning rest (Fig. S2C). Furthermore, ***α*** values were correlated between learning and post-learning rest (R = 0.71, 95% CI [0.32, 0.97]) and between pre- and post-learning rest (Fig. S2D), but showed only a non-significant trend between learning and pre-learning rest (R = 0.43, 95% CI [-0.04, 0.84]). Other PRG scaling exponents failed to show consistent behavioral-state-dependent changes (Fig. S1F, H, J), highlighting the potentially privileged role of ***α*** for characterizing the dynamical regime of neuronal networks. We emphasize that ***α*** represents a high-order network feature distinct from simpler metrics: for example, average firing rates, coefficient of variation (CV), pairwise correlations, synchrony, and excitatory/inhibitory ratios did not correlate with ***α*** during sleep/rest (Fig. S2E-H).

The state-dependent modulation of ***α*** suggests a link to the functional processes underlying memory. Given that post-learning rest is known to enhance subsequent recall (*1, 2*), we tested whether ***α*** during this period was related to memory retention during the post-probe session. Sessions with better memory retention exhibited lower ***α*** values, resulting in a significant negative correlation between normalized time at goal locations and ***α*** during post-learning rest (Fig. 1C-E, left panels). In contrast, ***α*** measured during post-learning rest did not correlate with performance during the pre-learning probe (R = 0.24, 95% CI [-0.18, 0.6]).

Because ***α*** values remained stable between pre- and post-learning rest (Fig. S2D), we examined whether the pre-learning rest dynamical regime alone could forecast future behavioral performance. Indeed, pre-learning rest ***α*** significantly predicted subsequent recall during the post-probe (Fig. 1D-E, right panels), but showed no relationship with baseline performance during the pre-probe (R = 0.24, 95% CI [-0.18, 0.6]). When incorporating ***α*** alongside other PRG parameters into a linear model to predict memory performance, ***α*** was the only significant contributor to the model’s predictive power (Fig. S3C). This predictive relationship between ***α*** and memory recall was robust across timescales ranging from 25 ms to 16 s (Fig. S4, S5), suggesting that prediction was also maintained in a scale-free regime.

Our results indicate that although ***α*** fluctuated across animals and recording days, it remained relatively stable from pre- to post-learning rest (Fig. 2G Fig. S2C-D); consequently, even pre-learning values predicted subsequent memory recall. The consistency of this relationship suggested that the predictive power of ***α*** might be invariant across animals. To test whether this is the case, we employed a leave-one-animal-out cross-validation procedure. For each animal, memory retention was predicted using a linear model trained exclusively on data from the remaining animals (a separate model was fitted for pre- and post-learning rest). The predicted values significantly correlated with actual memory retention for both pre- and post-learning rest (Fig. 1F, S3D, E). Notably, these correlations were significantly stronger than those obtained from control models where ***α*** values, training data, and retention scores were shuffled (Fig. 1F-G).

Consistent with previous reports (*15*), the expanded dataset, including newly acquired data, confirmed that the reactivation frequency correlated with memory retention at goal locations (Fig. 3A-B). We thus asked whether reactivation frequency and ***α*** were related. While we found significant correlation, with lower ***α*** values in post-learning rest associated with higher reactivation frequency (Fig. 3C), partial correlation analysis revealed that ***α*** remained a significant predictor of memory retention even when controlling for reactivation frequency (Fig. 3D). Conversely, reactivation frequency no longer predicted memory retention once its correlation with ***α*** was accounted for (Fig. 3E).

**Figure 3.**
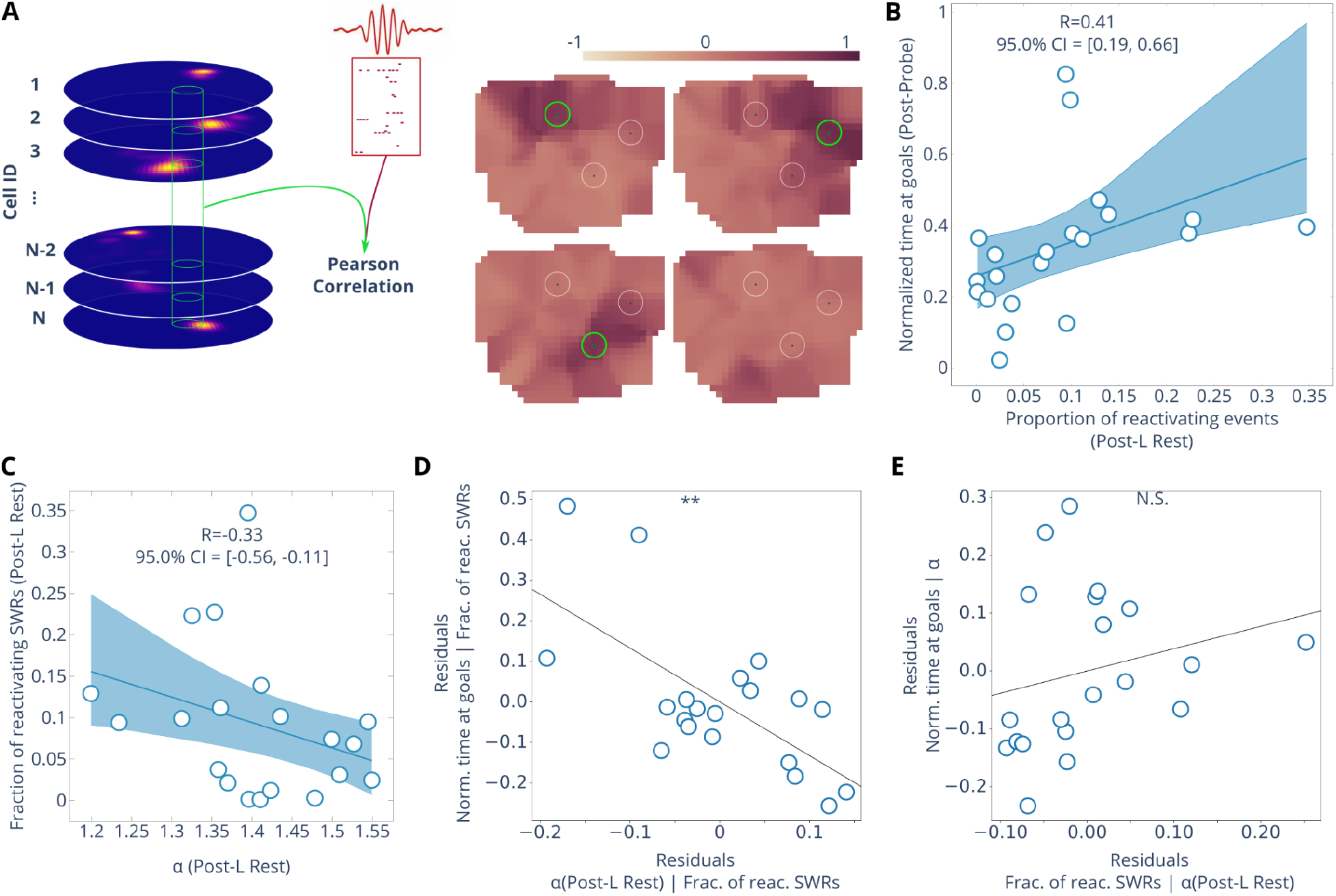
Prediction power of *α* is independent of reactivation. (**A**) Left: reactivation is assessed as Pearson correlation between population vectors (PV) from the place-rate map of place cells and SWR activity during Post-Learning Rest. Right: reactivation maps of four SWR events, pixel color indicates the Pearson correlation coefficient between the corresponding PV and SWR activity, green circles indicate reactivated goals, the SWR event at bottom right does not reactivate goals. (**B**) Correlation between the proportion of SWRs that reactivate goal zones and the normalized time spent around goals. (**C**) Correlation between ***α*** and the proportion of goal reactivation in post-learning rest. (**D**) Correlation residuals when correlating ***α*** and time at goals while controlling for the proportion of goal reactivation (partial correlation *r* =− 0. 66, *p* < 0. 01). (**E**) Correlation residuals when correlating the proportion of goal reactivation and time at goals while controlling for ***α*** (partial correlation *r* = 0. 23, *p* > 0. 1).

To further demonstrate that ***α*** predicts retention independently of reactivation, we recalculated ***α*** after excluding time windows containing sharp-wave ripples (SWRs) and high-synchrony periods. Even when restricted to these non-reactivation periods, ***α*** continued to correlate with memory recall (Fig. S6A). Together, these results suggest that while ***α*** identifies a dynamical regime that facilitates reactivation, it also regulates memory stabilization through mechanisms beyond ensemble reactivation. Consequently, we looked for further circuit factors that might influence ***α***, to identify additional mechanisms of memory stabilisation during offline periods.

While simple network activity statistics do not influence ***α*** (Fig. S2E-H), its value evidently depends on the underlying correlation structure, which could be constrained by the temporal firing characteristics of individual neurons. For instance, cells with very disparate intrinsic timescales (measured via the spike autocorrelation decay time) may fail to synchronize effectively; alternatively, cells with long intrinsic timescales could serve as “hub-cells,” weakly coordinating multiple fast-firing neurons that would otherwise remain uncorrelated. We found that sessions varied in the heterogeneity of these intrinsic timescales, specifically in the degree to which timescale distributions skewed toward longer constants, which we quantified by the Quartile Coefficient of Dispersion (QCD; Fig. 4A-B). The QCD correlated positively with ***α*** (Fig. 4C) and negatively with memory retention, suggesting that more uniform intrinsic timescales favor memory stabilization (Fig. 4D). However, this heterogeneity did not fully account for the relationship between ***α*** and memory retention: they remained significantly correlated even when controlling for QCD (Fig. 4E). However, the correlation between QCD and memory retention was no longer non-significant when accounting for ***α*** (Fig. 4F).

**Figure 4.**
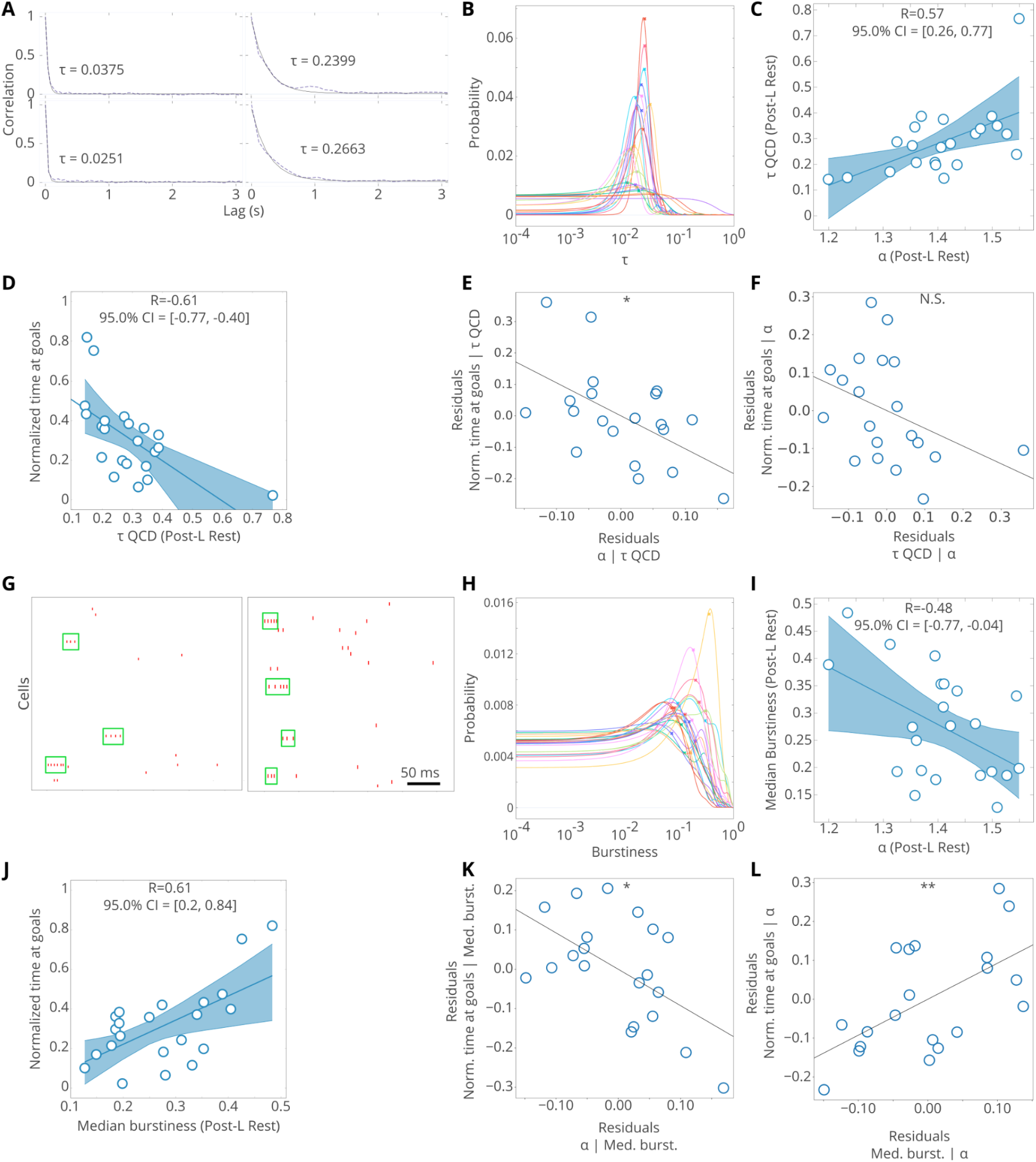
Distribution of cell activity parameters and scaling of activity variance. (**A**) Autocorrelation function of four cells and their intrinsic timescale (**τ**) measured by their time constant with an exponential function fit. (**B**) Distribution of **τ** during post-learning rest using a kernel density estimate. Colors: different recording days; filled circles: distribution median. (**C**) Correlation between quantile coefficient of dispersion (τ *QCD*) of **τ** distribution and ***α***. (**D**) Correlation between τ *QCD* during post-leaning rest and time at goals. (**E**) Residual plot for ***α*** and time at goals when controlled for τ *QCD* (partial correlation *r* =− 0. 54, *p* < 0. 05). (**F**) Residual plot for τ *QCD* and time at goals when controlled for ***α*** (partial correlation *r* =− 0. 38, *p* > 0. 1). (**G**) Rasterplot of bursting cells (green rectangles) from the two sessions. **(H)** Distribution of burstiness during post-learning rest. (**I**) correlation between median burstiness and ***α***. (**J**) Correlation between median burstiness and time at goals. (**K**) Residual plot for ***α*** and time at goals when controlled for median burstiness (partial correlation *r* =− 0. 56, *p* < 0. 05). (**L**) Residual plot for median burstiness and time at goals when controlled for ***α*** (partial correlation *r* = 0. 58, *p* < 0. 01).

Beyond intrinsic timescales, single-cell firing patterns may also influence the capacity for synaptic plasticity and thus directly impact memory retention. For instance, burst firing promotes plastic changes via dendritic calcium spikes (*28, 29*). We therefore examined whether the burst propensity of pyramidal cells during post-learning rest influences both ***α*** and memory retention (Fig. 4G-H). Interestingly, while burst propensity correlated more strongly with memory retention than with ***α*** (Fig. 4I-J), both measures remained independent predictors of memory retention when the other’s contribution was controlled for (Fig. 4K-L).

Finally, we asked how the composition of the analyzed network affects our conclusions. Many traditional predictors of memory, such as burst firing and reactivation frequency, focus exclusively on pyramidal cell activity. In contrast, we calculated ***α*** using both pyramidal cell and interneuron activity. When ***α*** was derived solely from pyramidal cells, it failed to predict recall performance (Fig. S7A-B). Notably, intrinsic timescale changes between pre- and post-learning rest were more positively skewed in interneurons than in pyramidal cells (Fig S7C). Consequently, the predictive power of ***α*** likely depends on the collective dynamics of the broader hippocampal circuit, including the interplay between excitatory and inhibitory populations.

## Discussion

Statistical physics-based approaches provide a unique lens for investigating how neural activity, captured at the level of individual spikes, impacts collective circuit regimes of operation that govern behavior. Using the PRG, we analyzed hippocampal neural activity during sleep/rest periods and its relationship with learning. Consistent with previous reports (*30*), we observed power-law scaling of multiple activity statistics across animals and sessions. While such scaling has traditionally been interpreted as a sign of closeness to criticality, a view that has not been without controversy (*31–33*), our analysis instead focused on the quantitative impact and predictive power of individual scaling exponents identified by the PRG. We found that the scaling exponent of variance, ***α***, evaluated during offline periods, strongly correlated with learning performance. Crucially, we demonstrated that ***α*** can serve as a transferable biomarker, because models trained on subsets of animals predicted memory performance of independent cohorts. Furthermore, ***α*** remained relatively stable across sessions and even predicted recall before learning took place (in pre-learning rest), suggesting that a sustained dynamical regime determined the capacity of hippocampal circuits for subsequent memory stabilization. While remaining relatively stable, nonetheless ***α*** exhibited behavioral-state-dependent shifts. This aligns with reports suggesting modulation of the proximity to criticality during the sleep-wake cycle (*22*). Other criticality-based parameters that measure instantaneous transitions also correlate with sensory discrimination (*21*), indicating that both fast and slow fluctuations in network dynamics can influence behavioral output.

While reactivation frequency correlated with both recall and ***α***, its predictive power vanished once its relationship with ***α*** was accounted for, suggesting that the broader network dynamical regime largely governs the rate and quality of memory replay. Memory stabilization requires the selective reactivation of specific memory traces (*26*) originating from either local circuit mechanisms or upstream inputs (*34*). Our results suggest that the low ***α*** dynamical regimes are a prerequisite for this process, either facilitating local selection or gating the effective expression of these upstream influences. Beyond selection, the dynamical regime could also determine coding efficiency (*35, 36*). Because lower ***α*** values correspond to more independent (i.e., less correlated) global activity, the lower ***α*** regimes we observed during successful recall might facilitate the formation of orthogonal CA1 assemblies that are more easily decoded by downstream targets (*37*). We further observed that ***α*** and burst propensity were correlated, yet both independently predicted retention. This indicated that scale-free dynamics reflect cellular-level features and, in addition, physiological affinity for dendritic-spike-mediated plasticity (*28, 29, 38*). In sum, our findings suggest that the optimal dynamical regime of hippocampal circuits modulates memory stabilization by enabling efficient reactivation and integration of memory traces, as well as the plasticity potential of the network

The low-***α*** regime was not a mere byproduct of pyramidal cell activity. While burst propensity contributed to memory retention, it did not fully account for the predictive influence of ***α***. Furthermore, the failure of pyramidal-only populations to predict behavior underscored the need to account for the full network activity to assess the dynamical regime. Interneurons likely influence the segregation of reactivated cell assemblies (*39*), while excitation-inhibition balance governs overarching dynamics and constrains neural firing rates (*40, 41*). In fact, we found ***α*** to be sensitive to the intrinsic timescales of individual neurons, and that a balanced distribution of these timescales favored the ***α***-mediated regimes that promote memory stabilization. Precise tuning of intrinsic timescales is thought to enable the optimal integration of information across multiple temporal scales (*42*), a process that likely facilitates the effective transfer of reactivated memory traces to downstream cortical areas (*4, 5*).

Taken together, our results established that scale-free hippocampal dynamics, distinguished by a simple-to-compute yet collective metric, constitute a powerful, cross-subject predictor of successful recall that transcends individual cellular variability. These results further imply that the ability to maintain, or to tune the system towards, an “optimal dynamical regime” may be a fundamental requirement for effective systems consolidation. Future work investigating how these global regimes evolve in aging or neurodegeneration could provide new avenues for identifying and potentially restoring memory function.

## Acknowledgments

We thank Jago Wallenschus for outstanding technical support and Alessandro Treves for the helpful discussions. AI tools (Google Gemini) have been used for finding relevant literature and for correcting and editing the text. This work was supported by the

European Research Council 281511 (JC)

ISTA Interdisciplinary Project Committee grant (JC, GT, PŽ, FL)

JC Austrian Science Fund(FWF) I3713 (JC)

Austrian Science Fund(FWF) Cluster of Excellence program “Neuronal Circuits in Health and Disease”. (JC, GT)

Austrian Science Fund (FWF), grant no. PT1013M03318 (FL)

Marie Sklodowska-Curie action, grant agreement No. 754411 and No. 101066790 (FL)

## Author contributions

Conceptualization: PŽ, FL, GT, JC

Data collection: PŽ,DD,CB,ST,JRW

Spike-sorting and data curation: PŽ, DD, CB, ST

Data analysis for the paper and figures: PŽ

Supervision: JC, GT

Writing – original draft: PŽ, FL, GT, JC

Writing – review & editing: PŽ, FL, GT, JC, DD, CB

## Competing interests

The authors declare that they have no competing interests.

## Data, code, and materials availability

Data related to waking activity from (*27*) is freely available. The rest of the data will be available and code related to the data analyses will be published on zenodo.org upon publication of the work.

## Methods

### Experimental procedures

#### Animals

All animals used in this study were Long-Evans rats, 10-20 weeks old, weighing 300 − 400g. In total, the analysis was performed on 11 animals. Parts of the dataset used were previously published, n=5 animals, n=12 recording days, (*8, 27*). New data included n=6 animals, n=10 recording days). In 3 animals, Channelrhodopsin ChR2 was expressed in the CA1 region, using Adenovirus AAV1 with mDLx promoter (*43*). However, there was no light stimulation in the recording days used for this study.

All experimental procedures involving the unpublished data were carried out in accordance with the Austrian Federal Law for experiments with live animals and under a project license approved by the Austrian Federal Science Ministry.

#### Maze Apparatus

We used the same “cheeseboard” maze as previously described (*8*). The maze is made of a circular PVC board with a diameter of 1.2m and a thickness of 2cm. The maze has 177 food wells drilled on the board in a regular grid with a spacing of 8cm between the centers of the wells. Alongside the maze was placed the start box of size 30×20cm with 50 – 60cm in height. The start box is not covered and has doors lower than the walls, allowing the animal to be tracked inside. A black curtain surrounded the entire apparatus.

#### Behavioral paradigm

The behavioral protocol used in this study has already been published (*8*). On each recording day, rats were trained to locate hidden food rewards at three wells. All recording days followed the same protocol, starting with a *pre-probe* session during which the rats were free to explore the maze without any food rewards for 5-25 minutes. The rats were then put into a separate sleep box for the 20-90-minute pre-learning rest session. Then, in the “*Learning*” session, food well locations changed relative to the previous recording day and rats needed to learn the new reward locations. While animals learned the locations in a few (3-5) trials, these new locations were reinforced in 20-50 trials. Following this, animals rested in the post-learning rest session (20-120 min, in 1 animal 480 min). Finally, rats’ memory recall was tested in the second unrewarded *post-probe* session. To prevent rats from using odor-guided navigation to locate food pellets, the maze was covered with a food powder made from the same pellets that were given to rats and the maze was randomly rotated.

At the beginning of every experiment, rats were pre-handled and habituated to both the experimenter and the recording environment. After the initial habituation period, implantation surgery was performed, followed by a recovery period of a week. Subsequently, rats were put on a food restriction protocol during which their weight was slowly reduced to the target value of 85 − 88% of the starting weight. During this period, rats were exposed to the maze and trained to locate three food pellets (20mg) and, after consuming them, returned to the start box. At every trial start, the start box door would open, allowing rats to come out onto the maze, then the doors would close and remain so until rats located all three food pellets. Upon locating and eating all the pellets, the start box door would open, allowing the rats to return. To motivate them to return, food pellets were also placed in the start box. This procedure was repeated for 10-20 trials per training day, depending on the motivation of the rats. The whole protocol was repeated for 4-6 days, until the rats performed the given task consistently and until they reached the target body weight.

#### Drive implantation and recovery

The rats were put under Isoflurane anesthesia (1−2%) and treated with analgesics throughout the procedure. The craniotomies were open and drives were implanted so that the tetrode bundles are positioned over the dorsal CA1 (2-4mm ML, -2.5-5mm AP). During implantation, the tips of the tetrodes were positioned superficially, 1 to 1.4mm from the surface of the brain. After surgery, rats were allowed at least a seven-day recovery period. After the recovery period, the tetrodes were slowly lowered towards the CA1 region of the hippocampus over the following few days. Neuropixel 2 probes were fixed and chronically implanted so that the innermost shank lies at coordinates ML: 2.6mm, AP: -4.0mm.

#### Extracellular Recordings

In n=10 animals, the recordings were performed using microdrives consisting of 16 or 32 independently movable tetrodes, while in one animal (n=3 sessions) a Neuropix 2.0 electrode was used. The tetrodes were made from four tungsten wires twisted and heated to bind together. The wire used was 10 or 12µm thick, while the tips were gold-plated to achieve a target impedance of 400 - 500kΩ. Each bundle of tetrodes consisted of 16 tetrodes implanted in the right hemisphere. Drives with 32 tetrodes had two bundles implanted bilaterally. On each recording day, tetrode positions were tuned to ensure that they remained within the dorsal CA1, using LFP features as visual guidance. All tetrode data were recorded using Axona or Intan acquisition systems for electrophysiological recordings, digitized at 20kS/s or 24kS/s. In Neuropixel recording, the data was digitized at 30 kS/s using a custom PXIe data acquisition card connected to a computer via a PXI chassis (NI 1071, National Instruments, Austin, TX), and OpenEphys software was used to write the data to disk. Recording sites with units and visible ripple oscillations were selected for the recordings.

#### Isolating single-unit activity

For the published data, manual spike clustering procedures are described in (*27, 39*). For the remaining data collected with tetrodes, individual spikes were detected as events whose amplitude crossed the threshold of five standard deviations over the mean amplitude of the signal. The spikes were assigned to the clusters using a combination of automatic clustering with Mountainsort (*44*) and manual curation with Phy (https://phy.readthedocs.io/en/latest/). For the Neuropixel recordings, data were sorted with Kilosort 4.0 (*45*). All putative units underwent manual curation in Phy2.

Units were classified as putative pyramidal cells or interneurons based on spike waveform features, autocorrelograms, and mean firing rates. Only recording days in which at least 32 cells were isolated were included in this data set.

### Position Recordings

On top of the microdrives were mounted LEDs that were used for position tracking. The position of the rat was recorded using a separate camera positioned above the maze. The camera was recording at 39.0625 frames per second and was synchronized with the acquisition system via a TTL pulse.

### Data analysis

Unless stated otherwise, data analysis pipelines are implemented using Linux core utils, Python programming language, and packages from its vast data analysis ecosystem: numpy (*46*), scikit-learn (*47*), scipy (*48*), statsmodels (https://www.statsmodels.org), pandas (https://doi.org/10.5281/zenodo.3509134), matplotlib (*49*), plotly (https://plotly.com/), seaborn (https://zenodo.org/records/12710), joblib (https://zenodo.org/records/16964648).

The quantities in the text are expressed as mean ± standard error, unless stated otherwise. In the text N.S. denotes p > 0.1, ∗ denotes p < 0.05, ∗∗ denotes p < 0.01, ∗ ∗ ∗ denotes p < 0.001, ∗ ∗ ∗∗ denotes p < 0.0001.

#### Bootstrapping correlations

Confidence intervals for correlations were computed using the bootstrap procedure by repeatedly drawing 5000 random samples (with replacement) from the original dataset, ensuring each “bootstrap sample” is the same size as the original. For each iteration, the Pearson correlation is recalculated. This generates an empirical distribution of the correlation coefficient, allowing us to determine confidence intervals.

#### Correlation between alpha and simple measures of activity (Fig. S4G-J)

Related to tables in figure S4G-J. All measurements were computed in pre- and post-learning rest sessions. All correlations were bootstrapped. Individual confidence intervals were set so that the overall significance level is 0.05, using Bonferroni correction. Data was binned using the same Geometric ISI strategy as for the other analysis.

- Mean cell firing rate: for each cell, the mean firing rate was computed in the given session and the mean across all cells was used;
- Mean pairwise corr.: mean of cross-correlations between binned activity traces, overall possible cells pairs;
- Population firing CV: coefficient of variation of population spike count
- Net. synchrony: mean synchrony in the network, where synchrony is defined as the fraction of pyramidal cells that are simultaneously active in a temporal bin;
- Net. synchrony CV - coefficient of variation of synchrony in the network;
- IE score: inhibition/excitation score is the difference between the average firing rate of interneurons and pyramidal cells, divided by the sum of the two.

#### Memory Retention Performance

Memory retention performance was evaluated as normalized time spent in reward zones, normalized relative to the baseline (Figure S4A, B). Reward zones were centered at the food well locations with a 12.5 cm radius. We first measured the time in the real reward zones *RZ*_*time*−*real*_ . Then we randomly sampled 1000 reward configurations outside of existing reward zones that the animal visited and calculated the time spent at the sampled rewards *RZ*_*time*−*sampled*_ . Finally, we calculated the following index:

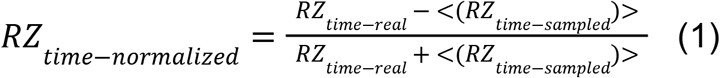

where the <… > denotes the average of given values. The normalized quantity assumes values from [− 1, 1], where value of 1 would occur if the animal spent all time in the reward zones, while a value equal to − 1 would occur if the animal never visited any of the reward zones. Memory retention was evaluated only during the first two minutes of Probe sessions. In all recording days used in the study *RZ*_*time*−*normalized*_ > 0.

#### Phenomenological renormalization group

Consider a population of N neurons, where the temporarily binned (details below) activity of the *i*-th neuron is denoted by *u*(*i*). Neurons are grouped during each step *k* of the phenomenological renormalization group (PRG) process based on the criterion of the maximum pairwise Pearson correlation coefficient. Denote by 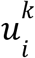 the activity of the coarse-grained unit *i* at step *k* of the PRG. Initially, the activities 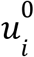 coincide with the original cell activities. Each step begins by selecting the most correlated pair of cells (*i, j*), and merging their activities into a new coarse-grained variable:

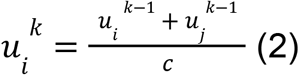

where *c* is such that the mean non-zero activity is equal to one, while zeros are preserved. Subsequently, we merge the next two most-correlated cells and continue until we form *N*/2 coarse-grained units, each comprising the aggregated activity of K=2 original cells. In the next iteration of the PRG (k = 2), each coarse-grained unit comprises *K* = 4 cells, for a total of *N*/*K* (*i. e. N*/4) coarse-grained units. This procedure is repeated iteratively until only one unit remains. At each PRG step, k, each unit comprises K=2^k^ original cells.

#### Temporal binning of data

The duration of temporal bins, δ*t*, used in the PRG analysis was calculated as the geometric mean of the inter-spike intervals, *d*_*i*_ (*i* = 1,…, *N*), across all cells:

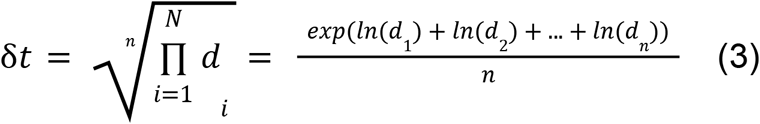

#### Pairwise correlations

Pairwise Pearson correlations were computed between units at the given level of PRG. If *u*_*i*_ represents the activity of the *i-*th unit, the Pearson correlation between units *i* and *j* was calculated as:

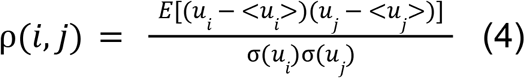

#### Activity variance and scaling exponent α

To compute the activity variance of each coarse-grained unit at every PRG step *k*, we first traced the *K* cells that comprise the given unit. Then we summed the activity of the traced cells and calculated the variance. We plotted the mean activity variance as a function of *K* on a logarithmic scale. The scaling exponent *α* is the slope of the line fitted to the log-log plot of the activity variance vs. K.

#### *Distribution of activity, free energy F, and scaling exponent* β

We denote with *Q*_*K*_ (*x*) the distribution of non-zero activity for the coarse-grained units comprising *K* = 2_*k*_ cells, corresponding to the PRG step *k*. Then, for a given coarse-grained unit, the probability of silence, *P*_0_, is estimated as the number of zero-bins divided by the total number of bins. The free energy, *F*, is defined as the negative logarithm of the probability of silence, i.e. *F* = − *ln*(*P*_0_). We plotted the mean free energy as a function of *K* on a log-log scale. The scaling exponent β is the slope of the line fitted to the log-log plot, i.e. *logF* = β · *logK* + *A*.

#### Spectrum of the covariance matrix and scaling exponent μ

For each PRG step *k*, we isolated the original cells that contributed to the each coarse-grained unit 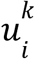. Among this group of cells, we calculated the covariance matrix 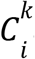 and its eigenvalues 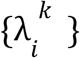. Each eigenvalue had an associated fractional rank defined as the eigenvalue rank divided by *K*, i.e.:

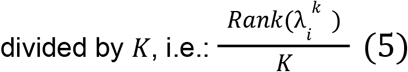

Eigenvalues were obtained for all units of the k-th PRG step, and the mean eigenvalues, λ, were plotted as a function of their fractional ranks. Then, the scaling exponent μ was obtained by fitting the following function to the eigenvalue-fractional rank relation:

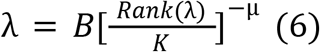

#### Autocorrelation time (intrinsic timescale) and scaling exponent z

The autocorrelation time, τ_*c*_, was calculated for every PRG step *k* and refers to the exponential decay constant of the population autocorrelation function, *C*(*t*).

We computed the autocorrelation function from the binned activity time series (bin duration estimated at k = 0). For each PRG step *k*, the binned activity of each unit was z-scored, and the normalized autocorrelation, *C*(*t*), was computed:

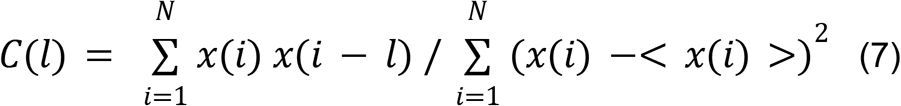

Where *N* denotes the number of elements in *x* and -*N* ≤ *l* ≤ *N* is the autocorrelation lag. Finally, the average autocorrelation over all units was calculated. The decay constant τ_*c*_ (*K*) was obtained by fitting an exponential function of the form:

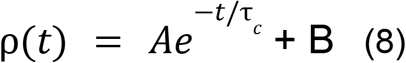

to C(t), where *t* = *l* · *bin length* represents time. The scaling exponent *z* is the slope of the line fitted to the relationship between *log*[τ_*c*_ (*K*)] and *log K* .

#### Shuffling and simulation procedures for null model PRG exponents

To disrupt interactions between cells while maintaining the firing properties of single cells, we performed two shuffling/randomization and one simulation procedure.

In the first procedure (shuffle, shuffle-10, shuffle-100, etc), we shifted the binned spike train of each cell independently of other cells. For each cell, a shift amount *N*_*shift*_ was randomly chosen and each bin was moved forward by *N*_*shift*_ bins, while the bins from the end were moved to the beginning of the binned spike train. Repeating this procedure independently for each cell disrupts correlations between cells, while preserving the firing rate of each individual cell. We performed this procedure for the whole session (“Shuffle”), as well as in shorter sliding windows, with *N*_*shift*_ between 10 and 10000 bins (Sh-10, Sh-100, Sh-1000, Sh-10000). For each group and each recording day, we performed 100 independent shuffling of the spike trains.

In the second procedure, we generated surrogate data by matching firing rates of individual cells on sliding windows of varying sizes. For each recorded cell, we randomly drew a binned spike train from a multinomial distribution with parameters set in such a way to match each cell’s firing probability in the sliding window. Repeating this procedure independently for each cell disrupts correlations between cells, while preserving the firing rate of each individual cell. We performed this procedure for the whole session (“Random”), as well as in shorter sliding windows of 10, 100, 1000 and 10000 bins (R-10, R-100, R-1000, R-10000). For each group, each recording day, and randomization setting, we performed 100 random draws.

Finally, for each recorded cell, we simulated spike trains by randomly sampling from a Poisson distribution with the rate parameter set to the cell’s firing rate (Ind. Poisson). Repeating this procedure independently for each cell disrupts correlations between cells, while preserving firing rate of each individual cell. We performed this procedure for the whole session (“Ind. Poisson”) by performing 100 random draws for each recording day.

#### Predicting memory performance across animals

To investigate whether the scaling exponent *α* represents a quantity that is transferable between animals, we performed cross-validation by excluding each animal from the training set and using the excluded animal for testing.

Based on our dataset of 11 animals, we generated 11 train-test dataset splits. The test part of each split consisted of all days recorded from the excluded animal, while the training part consisted of all remaining sessions. On each split, we trained a linear regression model that predicted normalized time around goals based on the scaling exponent *α*. We then tested models on test splits that consisted of recording days from animals that the models did not see during training.

We evaluated the prediction quality in two ways. First, for excluded animals, we correlated the predicted normalized time around goals with the measured normalized time around goals. Second, we calculated the coefficient of determination of predicted normalized time around goals (*R*^2^) and compared it with the shuffled distribution. We generated a shuffle distribution by randomizing labels (norm. time at goals) when training models (200 random draws).

#### SWR and HSE detection

We detected high synchrony events (HSEs) using a method very similar to that used in (*50*), based on the combined multiunit activity (MUA) of the principal cells. First, MUA was binned into 1.5 ms bins and smoothed with a Gaussian kernel with SD = 15 ms. Next, we z-scored the smoothed MUA and marked all continuous bin blocks with z-score > 3 as HSEs. We determined the beginning of HSE by extending left from the peak while the z-scored MUA was greater than 1. We determined the end of HSE by extending right from the peak while the z-scored MUA was greater than 1. All neighbouring HSEs that occur within 50 ms were merged. We kept only HSEs with a duration between 50 ms and 1000 ms. We kept only HSEs with at least 5 emitted spikes and a number of active cells larger than 4 or 10% of the total number of cells.

To detect SWRs we analyzed the local field potentials of tetrodes located in CA1. For each tetrode, we first performed a band-pass filter to 150-250 Hz and calculated the instantaneous power of the signal. Then, we summed the power from the signal of all CA1 tetrodes and z-scored it. The SWR peaks were then identified as moments in which the z-scored power was higher than 6. To determine the beginning and end of an SWR event, we extended from the peak until the z-scored power was larger than 1.2.

Finally, for the analysis related to Figure S6A, we removed periods that corresponded to both SWRs and HSEs.

#### Place field calculations

#### Occupancy Maps

Estimates of the probability of visiting each part of the maze are represented as occupancy maps. To compute them, we assigned the recorded *x* − *y* positions to the 3 × 3 *cm* ^2^ spatial bins and then smoothed the bin counts with a Gaussian kernel with *SD* = 2.

#### Firing Rate Maps

Spatial firing rate maps were calculated for every cell separately. First, we divided the whole maze into 3 × 3 *cm*^2^ spatial bins. We next counted the number of emitted spikes in every spatial bin and smoothed the bin counts with a Gaussian kernel of *SD* = 2. Then, we divided smoothed bin counts with the smoothed occupancy map to compute estimates of firing rates for every position bin. We used only periods of mobility, with *speed* > 3 *cm*/*s*.

#### Spatial Information Content

We calculated the spatial information content of the principal cells in bits / spike as in (*51*) :

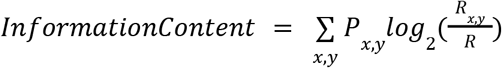

where *P*_*x,y*_ is the occupancy of the bin (*x, y*), *R*_*x,y*_ is the mean firing rate of the cell in the bin (*x, y*) and *R* is the overall mean firing rate of the cell.

#### Spatial Information Criteria

In the reactivation analysis, we used only cells with spatial tuning. To determine if a cell had spatial tuning, we first created a per-cell shuffled distribution by randomly shifting the cell’s firing rate map along the x and y axes 200 times. For every random shift, we calculated the spatial information content to obtain a shuffle distribution. A cell had spatial tuning if its information content was greater than both the 95*th* percentile of the shuffled distribution and greater than the threshold of 2 bits/spike.

#### Population Vectors

For every position bin, a list of firing rates of all cells in that position bin is a population vector. To calculate population vectors, we stacked firing rate maps of analyzed cells along the z-axis and took the firing rates along all the spatial coordinates *x* − *y*.

#### Reactivation

Reactivation analysis was performed during the sleep/rest session *SS* (pre- and post-learning rest) and based on awake activity recorded during the first two minutes of the post-probe session. First, population vectors (PVs) were calculated based on the first two minutes of the post-probe session. For each of the three goals, a list of *PVs* was compiled from the 12.5 cm reward zone and only PVs from these lists were used for the reactivation analysis. The activity corresponding to SWR events was then isolated during post-learning rest. Each candidate event had an associated activity vector with cell firing rates. Activity during all candidate events was compared with PVs of all three goals, allowing a detailed look at per-goal reactivation. For a given goal *G*_*i*_ and a given candidate event *e*, the Pearson correlation coefficient was calculated between the activity vector of the event and all PVs related to the goal *G*_*i*_ . The maximum correlation coefficient ρ_*max*_ was selected.

Next, shuffle *PVs* were generated by randomly shifting the awake firing rate maps of each cell independently, along both axes. Values that shifted beyond the last position were reintroduced at the first. This procedure maintains single-cell place selectivity but breaks spatial correlations among cells. Random shuffling was performed 200 times and for each iteration, the maximum correlation ρ_*max*_ between the shuffled PVs and SWR activity was established to generate the shuffled distribution of ρ_*max*_ .

If the original ρ_*max*_ correlation was significant (*p* < 0. 05), it was compared with the shuffle distribution. If ρ_*max*_ exceeded the 95th percentile of the shuffle distribution, the candidate event *e* was marked as a significant reactivation event for the goal *G*_*i*_ . This procedure was repeated for the two remaining goals, allowing each candidate event to reactivate more than one goal at a time.

To investigate the relationship between reactivation and memory retention, for each recording day, a fraction of significant reactivation events was calculated. This was correlated with the normalized time spent around the goal. All correlations were bootstrapped 5000 times.

Reactivation maps displayed in Fig. 3A were calculated for each event separately. Each PV, from the whole map and not only the reward zones, was correlated with the activity vector of the event. The reactivation map is a 2D array of Pearson correlation coefficients between PVs and the event’s activity vector.

#### Single-cell characteristic timescales

The characteristic timescale τ of a cell is defined as the exponential decay constant of the cell’s autocorrelation function. It was calculated for each cell, as described for the PRG scaling exponent *z* calculation above.

#### Burstiness

Cell burstiness was calculated on a single-cell basis. It is defined as the fraction of inter-spike intervals shorter than 10ms.

#### Kernel density estimation

To estimate the probability density functions on Figure 4B and I, we used Gaussian kernel density estimation. Bandwidth was calculated by multiplying the covariance matrix by the bandwidth factor *N*^−1/5^ calculated using Scott’s rule.

## Supplementary Figures

**Figure S1.**
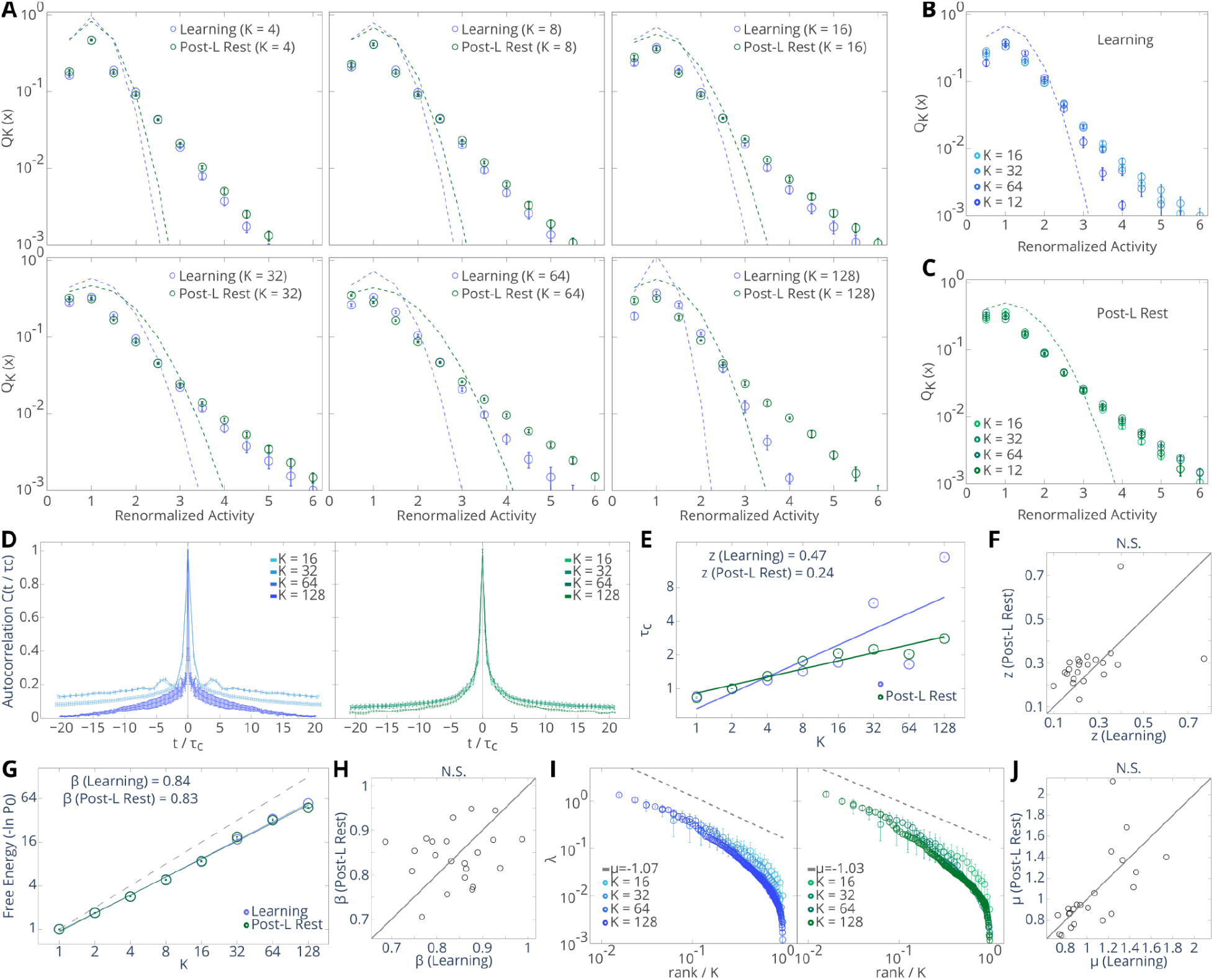
Other parameters of the PRG. See Methods for further details. (**A**) Distribution of normalised, non-zero coarse-grained activity for different PRG steps, K. Coarse-grained activity is renormalized at each step so that the mean of non-zero temporal bins is equal to 1. bin width: 0.5. Only activity in the range [0, 6] is shown. Data is pooled from all recording days. Circles indicate mean probability. Error bars (SEM) are smaller than the symbol size. The dashed lines are normal distributions with the mean and variance of the non-zero coarse-grained activity. (**B, C**) Coarse-grained activity distributions during learning and post-learning rest. (**D**) - autocorrelation function rescaled with autocorrelation time constant **τ**_**c**_, learning (left) and post-learning rest (right), averaged over recording days. (**E**) Scaling of **τ**_**c**_ in learning and post-learning rest (scaling exponent *z)*. (**F**) Exponent z during learning vs post-learning rest. (**G**) Scaling of free energy, F∝K^β^, during learning and post-learning rest. Data pooled from all recording days. Error bars are smaller than the symbol size. The dashed line denotes scaling with β = 1, expected for independent variables. (**H**) Exponent β in learning and post-learning rest. (**I**) Scaling of the eigenvalue spectrum for the covariance matrix of clustered activity during learning (left) and post-learning rest (right) is independent of K. Eigenspectra as a function of the fractional rank (rank/K) collapse onto the same power-law with exponent μ. Data pooled from all recording days. Circles indicate the mean eigenvalue λ. Error bars are inside the circles. Dashed gray lines: power law with the exponent μ (average across K). (**J**) Scaling exponent μ during learning and post-learning rest.

**Figure S2.**
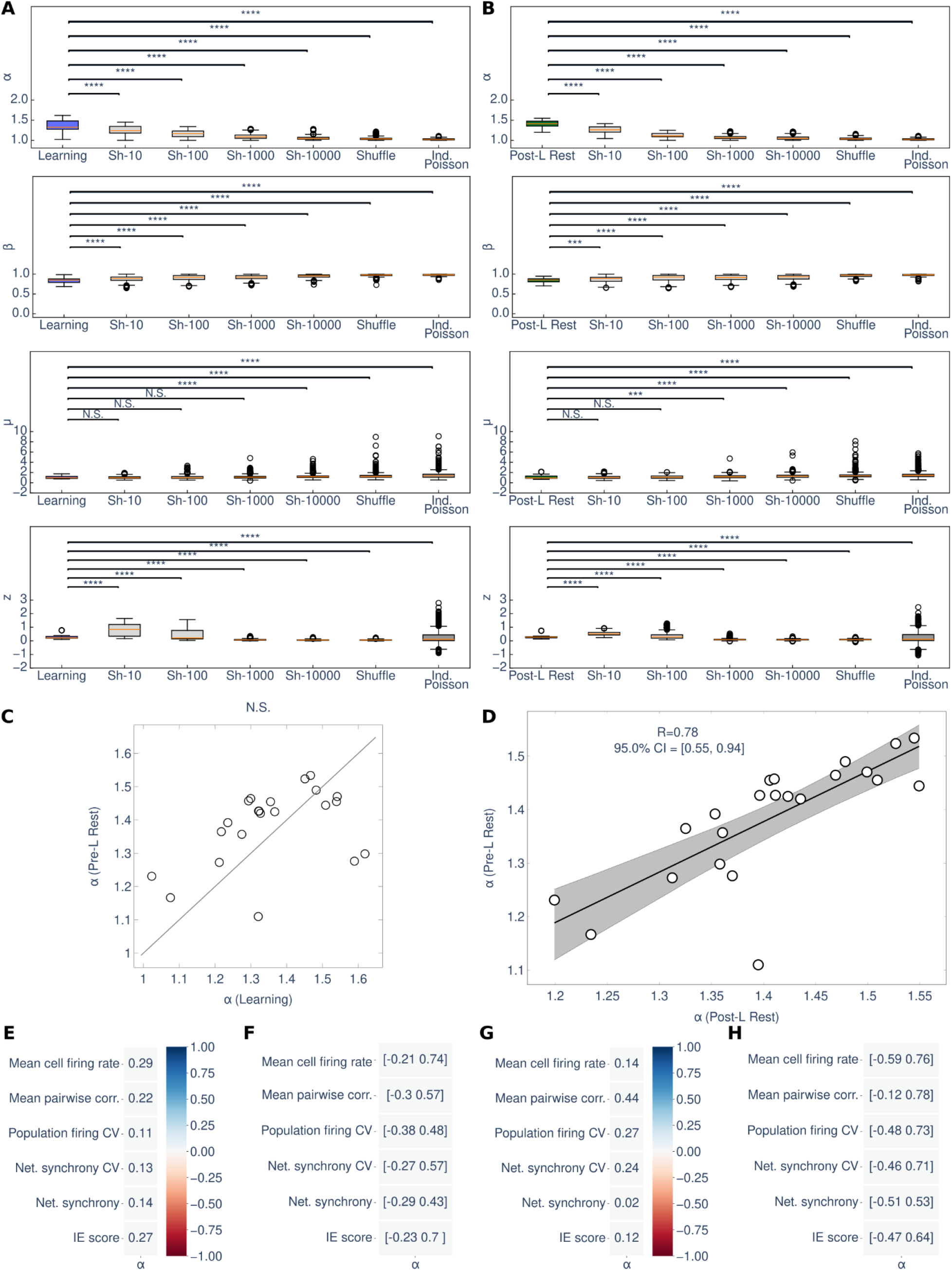
PRG scaling exponents in real and surrogate data. Sh-10: shuffling within a window of 10 bins, sh-100: 100 bins, etc. Shuffle: shuffling within the whole session, Ind. Poisson: data simulated from independently firing Poisson units. (**A**) Learning. (**B**) ost-learning rest. (**C**) Scaling exponent ***α*** in the learning and pre-learning rest (p>0.1, Wilcoxon signed-rank test). (**D**) Scaling exponent ***α*** in pre-learning rest is correlated with that in post-learning rest. (**E**) Correlations between *α* and several measures of network activity are not significant during pre-learning rest. (**F**) 95% CI for correlations of panel (E), (**G, H)** Same as (E, F), but for post-learning rest. N.S. p > 0.1, ∗ ∗ ∗ p < 0.001, ∗ ∗ ∗∗ p < 0.0001.

**Figure S3.**
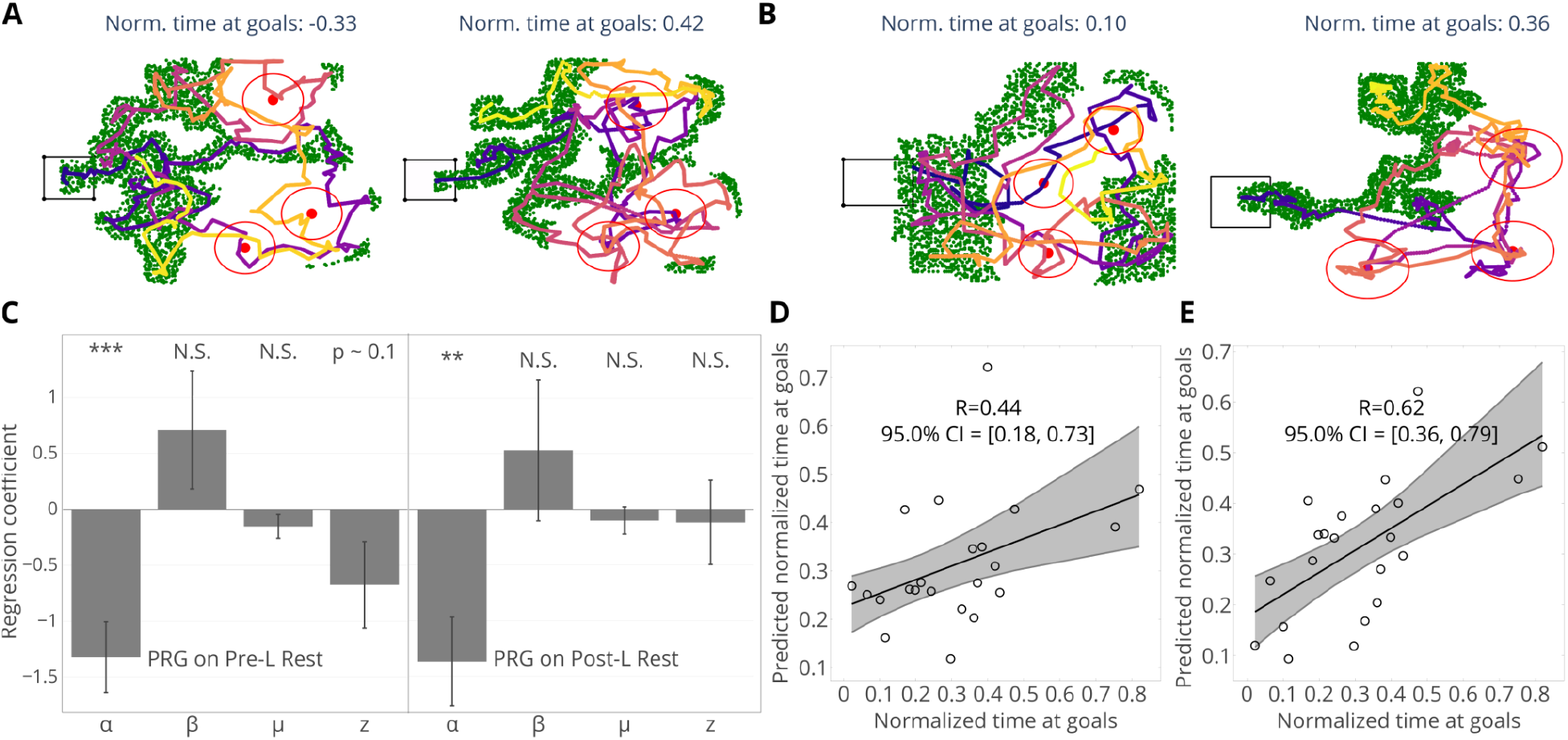
PRG exponents and memory recall. (**A**) sample trajectories on Pre-Probe (left) and Post-Probe (right) for the example session in Fig. 1. Black rectangle represents the startbox, red circles indicate goals, green dots indicate randomly sampled locations outside goal zones. (**B**) Sample Post-Probe trajectories from Fig. 2 with overlaid randomly sampled locations. (**C**) Regression coefficients from a multiple regression model of PRG scaling exponents on pre-learning rest (left) and post-learning rest (right). (**D, E**) Correlation between real and predicted time at goals using the “cross-animal” model trained on pre-learning rest (D) and post-learning rest (E) (same as Fig. 2 E,F but showing the bootstrapped correlations. N.S. p > 0.1, ∗∗ p < 0.01, ∗ ∗ ∗ p < 0.001.

**Figure S4.**
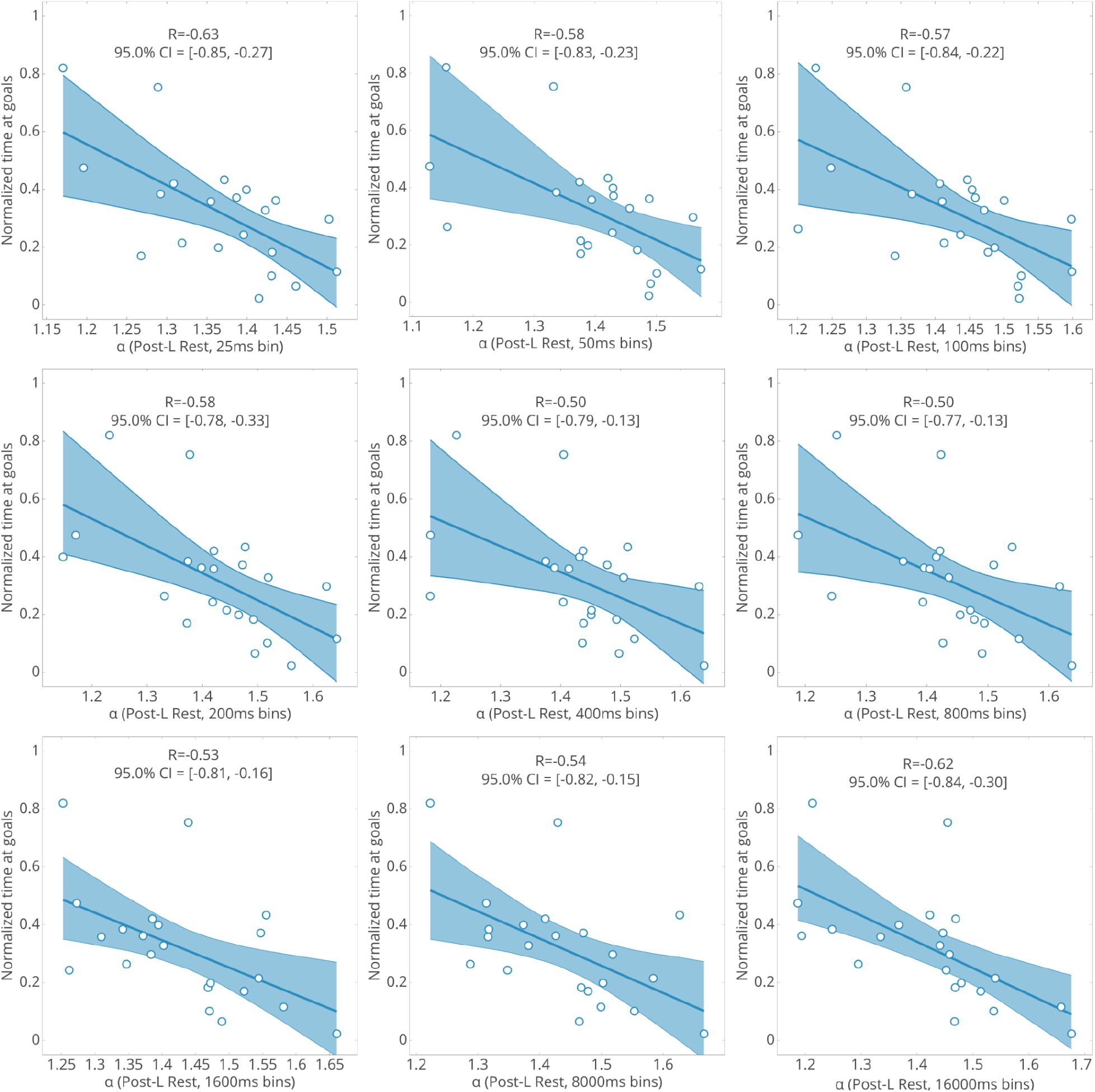
*α* - recall correlations over multiple temporal binning scales in post-learning rest. Scaling exponent ***α*** during post-learning rest correlates with the normalized time around goals during Post-Probe when ***α*** was calculated for a range of temporal bin durations.

**Figure S5.**
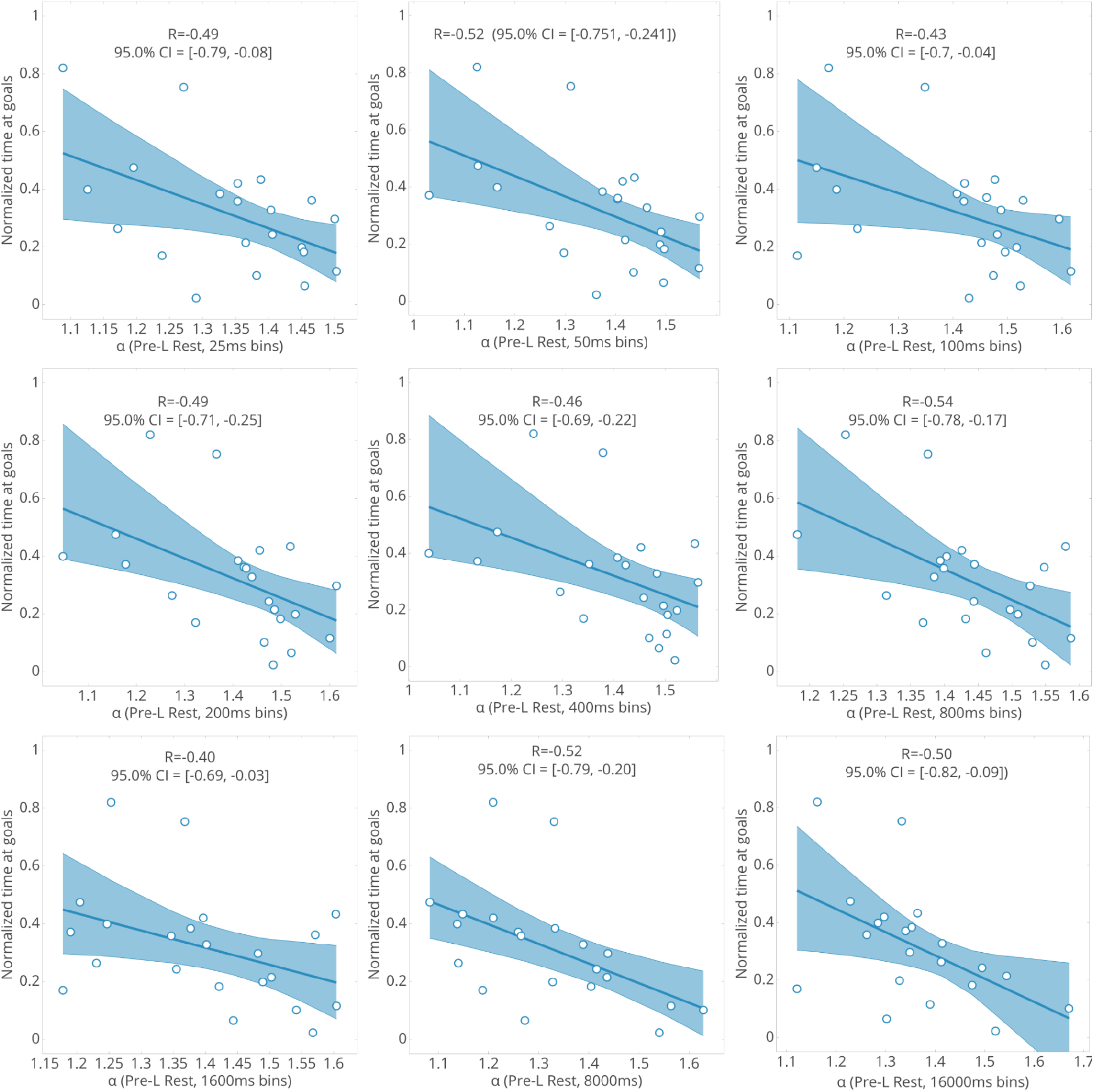
*α* - recall correlations over multiple temporal binning scales in pre-learning rest. Scaling exponent ***α*** during pre-learning rest correlates with the normalized time around goals during Post-Probe when ***α*** was calculated for a range of temporal bin durations.

**Figure S6.**
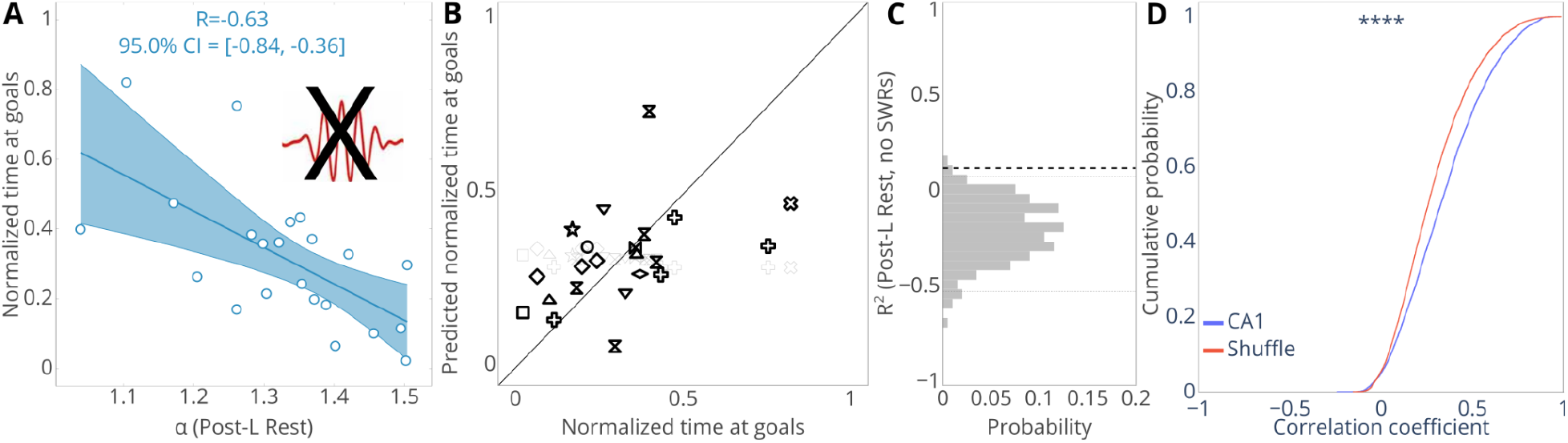
**(A-C**) Excluding SWRs does not affect the predictive power of the *α*. **(A)** Correlation between the ***α*** during post-learning rest when SWRs are excluded and the time at goals during post-probe. (**B**) prediction of memory retention using the cross-animal model when SWR periods are excluded during ***α*** calculation, every shape corresponds to a different predicted animal, light grey: predictions from a model trained on shuffled data. (**C**) The coefficient of determination of prediction shown in (B) grey histogram denotes the shuffle distribution, the thin lines show 95% CI, dashed black line denotes R^2^ observed on the data. (**D**) cumulative distribution function of Pearson correlation between goal population vectors and SWRs during post-learning rest, compared to shuffle (*p* < 0. 0001, Kolmogorov-Smirnov test), related to Fig. 3.

**Figure S7.**
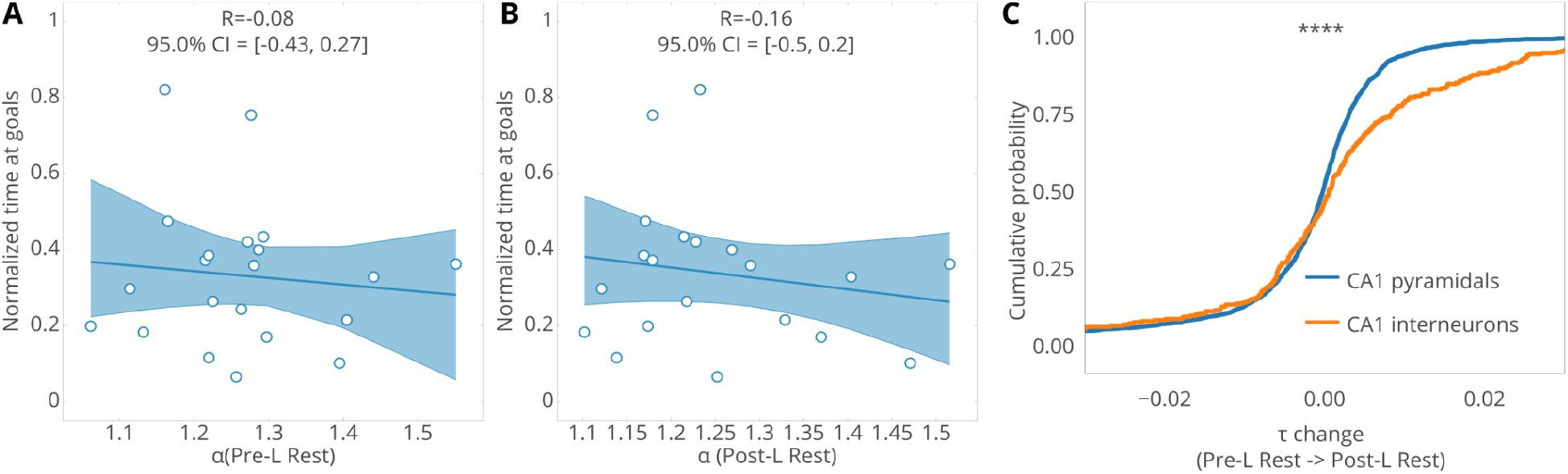
**(A,B)** Excluding interneurons breaks the correlation between the scaling exponent ***α*** and normalized time around goals (**A**) during pre-learning rest. (**B**) during post-learning rest. (**C**) cumulative distribution intrinsic timescale changes between pre-learning rest and post-learning rest is different between interneurons and pyramidal cells (p < 0.0001, KS-test) due to the positive skew in interneurons, albeit median change for fpr both cell types are close to zero, albeit significant (pyramidal cells: − 0. 00009 *p* < 0. 05, median change for interneurons: 0. 00084, *p* < 0. 05, Wilcoxon test)

## Notes

### Competing Interest Statement

The authors have declared no competing interest.

### Summary of Updates

Expanded abstract, expanded references, fixed typos, Figure 1A revised, slight changes in formatting.

